# Epithelial-mesenchymal plasticity: How have quantitative mathematical models helped improve our understanding?

**DOI:** 10.1101/122036

**Authors:** Mohit Kumar Jolly, Satyendra C Tripathi, Jason A Somarelli, Samir M Hanash, Herbert Levine

**Author notes:** Email: Herbert Levine.

## Abstract

Phenotypic plasticity, the ability of cells to reversibly alter their phenotypes in response to signals, presents a significant clinical challenge to treating solid tumors. Tumor cells utilize phenotypic plasticity to evade therapies, metastasize, and colonize distant organs. As a result, phenotypic plasticity can accelerate tumor progression. A well-studied example of phenotypic plasticity is the bidirectional conversions among epithelial, mesenchymal, and hybrid epithelial/mesenchymal phenotype(s). These conversions can alter a repertoire of cellular traits associated with multiple hallmarks of cancer, such as metabolism, immune-evasion, and invasion and metastasis. To tackle the complexity and heterogeneity of these transitions, mathematical models have been developed that seek to capture the experimentally-verified molecular mechanisms and act as ‘ hypothesis-generating machines’. Here, we discuss how these quantitative mathematical models have helped us explain existing experimental data, guided further experiments, and provided an improved conceptual framework for understanding how multiple intracellular and extracellular signals can drive epithelial-mesenchymal plasticity at both the single-cell and population levels. We also discuss the implications of this plasticity in driving multiple aggressive facets of tumor progression.

## Introduction

A remarkable feature that cancer cells use to evade therapy, metastasize, and drive tumor progression is phenotypic plasticity, i.e. ability of cells to switch back and forth among multiple phenotypes in response to varied internal or external signals (Hölzel et al., 2012). Plasticity is usually tightly controlled during adult homeostasis. It comes into play only when needed, such as during tissue repair, when resident stem cells give rise to cells that need to be replenished. However, during tumor progression, many of the molecular brakes against phenotypic plasticity are deregulated, enabling cancer cells to behave as ‘moving targets’ that can play ‘hide-and-seek’ with multiple therapeutic regimes (Roesch, 2015; Varga et al., 2014). In addition, these phenotypic conversions can facilitate adaptation by enabling genetically-identical cells to exhibit a diverse set of phenotypes and may also help fuel genetic evolution of cancer cells (Brooks et al., 2015; Mooney et al., 2016; Yadav et al., 2016).

A canonical example of such phenotypic plasticity that contributes significantly to both metastasis and drug resistance is epithelial-mesenchymal plasticity, i.e. the ability of cells to undergo a partial or full Epithelial-Mesenchymal Transition (EMT) and its reverse Mesenchymal-Epithelial Transition (MET) (Diepenbruck and Christofori, 2016). Interestingly, emerging evidence strongly suggests that these transitions are rarely ‘all-or-none’. Rather, cancer cells can often display a hybrid epithelial/mesenchymal (E/M) phenotype by combining various epithelial and mesenchymal morphological and/or molecular features (Jolly et al., 2015; Nieto, 2013; Nieto et al., 2016). Cells in this (these) hybrid state(s) can be much more tumorigenic and drug resistant as compared to those that are more fixed in an strongly epithelial or mesenchymal state (Biddle et al., 2016; Grosse-Wilde et al., 2015; Jolly et al., 2015). Thus, elucidating the underlying principles of these dynamic transitions is of foundational importance for countering the yet insuperable clinical aspects of cancer – metastasis and drug resistance.

Recent progress in dissecting the molecular mechanisms underlying these phenotypic transitions has enabled the development of quantitative mathematical models that can be used as hypothesis-generating tools to guide further experiments. In this review, we highlight how an integrative theoretical-experimental approach has helped us better characterize epithelial-mesenchymal plasticity. For instance, mathematical models capturing the dynamics of core EMT signaling network have predicted that cells can maintain a hybrid E/M phenotype stably, and that isogenic populations (cell lines) can contain admixtures of epithelial, hybrid E/M and mesenchymal subpopulations. These predictions have been validated by experimental observations showing different cell lines can contain subpopulations of different phenotypes in varying ratios.

### Why develop quantitative mathematical models?

Quantitative mathematical models offer us a powerful conceptual framework to elucidate underlying biological mechanisms and to propose new sets of experiments by generating falsifiable hypotheses. They can help interpret or explain the existing experimental data, confirm or reject alternate hypotheses, predict cellular behavior, and eventually guide further experiments (Mobius and Laan, 2015). They can decode the emergent dynamics of various regulatory networks and biological phenomena, and enable the experimental biologists to think more quantitatively in terms of regulatory dynamics. Mathematical models can also help unravel the principles that govern cancer progression, from the molecular scale all the way to the population level (Anderson and Quaranta, 2008). Thus, these models can aid in guiding optimal treatment modalities and can contribute to improved risk prognoses (Altrock et al., 2015).

### What is a quantitative mathematical model?

A model of any system is a replica that captures the system’s essential features and can thus be used to predict how the ‘original’ system would behave in a variety of conditions. Each model has its own assumptions, strengths and limitations, and is therefore suitable to answer a specific set of questions. In biology, we often use various pre-clinical models (cell lines, mouse models, patient-derived xenografts etc.) to investigate different phenomenon relevant to human biology, with an implicit expectation that lessons learned in these preclinical models can provide useful insights into the functioning of the human system. Broadly speaking, these biological models can be *in vitro* or *in vivo*. Similar to these model systems, a quantitative mathematical model is an *in silico* representation of the ‘original’ system, where a set of equations captures the essence of biological phenomenon through terms representing different objects involved in a phenomenon and interactions among them that govern that phenomenon (Figure 1A). A bidirectional communication among mathematical and experimental biologists has been fruitful in teasing out the mechanistic aspects of many biological processes such as timing and patterning of developmental events (Lewis, 2008; Oates et al., 2009; Shaya and Sprinzak, 2011).

**Figure 1:**
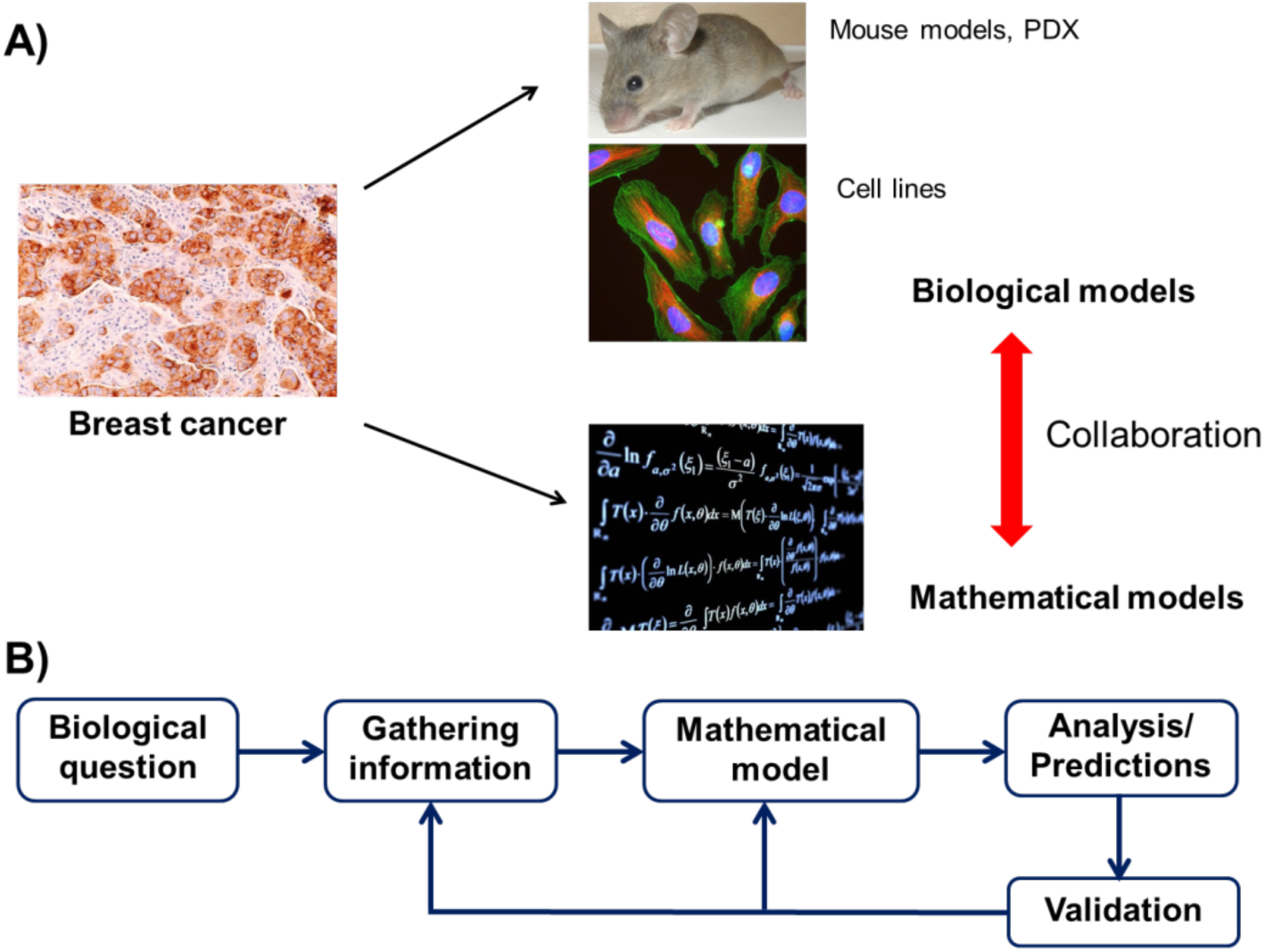
Introduction to quantitative mathematical models. **A**) Similar to biological models (e.g. cell lines, mouse models and, PDXs), mathematical models can capture certain aspects of tumor progression. Insights gained using both classes of models can be more than useful than through any one class solitarily. **B**) The process of developing, calibrating, and validating a mathematical model for a specific biological question. Generating predictions that can guide further experiments is the keystone of this integrative theoretical-experimental approach.

Just like biological models, mathematical models differ in scope and purpose (Mobius and Laan, 2015). For instance, different mathematical models developed to understand epithelial-mesenchymal plasticity have focused on different questions – (a) how do a set of transcription factors and microRNAs regulate the intracellular dynamics of a partial or full EMT/MET and modulate phenotypic heterogeneity in an isogenic population (Lu et al., 2013; Steinway et al., 2014; Tian et al., 2013)?; (b) how does cell-cell communication affect the spatial arrangement of epithelial, mesenchymal and hybrid epithelial/ mesenchymal cells (Boareto et al., 2016)?; and (c) how do cells alter their morphological and motility traits during EMT? As one may suspect, developing mathematical models to answer each of these questions requires quite different experimental data. Therefore, often times the scope of the model is decided by the data that are available – for example, whether longitudinal data are available either in discrete time points or in a more continuous fashion, whether data are available at a population-vs. single-cell level, whether the available data are merely for altered protein and transcript levels or they also includes morphology and motility aspects too etc. In this review, we will focus on a set of mathematical models that can be compared extensively against the existing experimental data.

### How does one develop a quantitative mathematical model?

As discussed earlier, the first step in developing a mathematical model entails being clear both about the biological question that the model should be able to answer, and the experimental data available with which to construct, calibrate and compare the model. Second, one must realize the implicit assumptions of different modeling frameworks and decide whether operating under those assumptions enables a reasonable replica of the ‘original’ biological system. These assumptions should always be judged in the light of the question/phenomenon of interest. Third, one should strive to accurately incorporate multiple key features of a phenomenon in one’s model. Finally, the model should be validated by comparing the predictions of the model in cases where robust experimental data are available *a priori*. Subsequent to model validation, one can generate predictions that can be tested experimentally and confirmed or falsified (Figure 1B).

Generating falsifiable predictions is the most useful application of developing mathematical models. Therefore, simply fitting experimental data to a model does little to contribute to new knowledge. Rather, one should seek to stick the model’s neck out after it is fitted and try to falsify it’ (Gunawardena, 2014) by predicting how the ‘original’ system (often, the biological model system being studied) would behave under altered conditions, such as by introducing genetic mutations or overexpressing a specific gene.

What happens if there is a mismatch between the prediction of the mathematical model and the experimental results generated? This mismatch can occur due to multiple reasons, such as (a) underlying assumptions of the model are not entirely valid, (b) the model is not robust, i.e. relatively small changes to the model or its parameters dramatically change the behavior of the model, and/or (c) technical inaccuracies in running experiments and/or model simulations. Once the underlying reason(s) is (are) identified, and predictions of the mathematical model score well with experimental results, this iterative cycle can continue to identify the next set of exciting research directions to be answered using the same or a different mathematical and/or biological model(s), as applicable.

### How can epithelial-mesenchymal plasticity be represented by a set of mathematical equations?

An exemplary biological phenomenon in which mathematical modeling has helped provide useful biological insights is that of epithelial-mesenchymal plasticity. This plasticity arises via a gene regulatory network that controls reversible switches between phenotypes, and has implications for numerous key biological processes in normal and disease states. For example, in the context of cancer, phenotypic switching between epithelial and mesenchymal states via EMT and MET drives cancer progression, metastasis, and therapy resistance. These epithelial and mesenchymal cells have distinct morphological and molecular features. For instance, epithelial cells have E-cadherin (CDH1) localized at the cell membrane, which contributes to adherens junctions. Conversely, mesenchymal cells lack E-cadherin and typically have higher levels of Vimentin (VIM), N-cadherin (CDH2), and αSMA (smooth muscle actin). Thus, EMT and MET typically involve widespread changes in gene expression, microRNAs, and epigenetic profiles, as well as cytoskeletal reprogramming (De Craene and Berx, 2013). An understanding of the set of molecular players of interest and the interactions among them can facilitate development of a mathematical model that can trace these changes during EMT and MET, and potentially highlight novel areas of susceptibility to therapeutic targeting.

The first set of mathematical models developed for EMT/MET focused on a specific question: can the underlying EMT/MET regulatory network enable the existence of a stable hybrid epithelial/mesenchymal (E/M) phenotype; and if so, what is the molecular signature of this hybrid E/M phenotype (Lu et al., 2013; Tian et al., 2013)? These efforts at addressing this question modeled the interactions among two sets of microRNAs and two sets of transcription factors that were reported to govern EMT/MET in multiple cell lines – miR-34, miR-200, ZEB, and SNAIL (Bracken et al., 2008; Gregory et al., 2008; Kim et al., 2011; Park et al., 2008; Siemens et al., 2011) (Figure 2A). The models predicted that under certain conditions a hybrid E/M phenotype can be stable, and that in isogenic populations multiple phenotypes can co-exist, including {E/M, M}, {E, M}, and {E, E/M, M} (Figure 2B). These predictions were later validated by experiments demonstrating subpopulations of E, hybrid E/M, and M phenotypes in varying ratios in cell lines across multiple cancer types, as assessed by flow cytometry and immunofluorescence (Andriani et al., 2016; Grosse-Wilde et al., 2015; Jolly et al., 2016; Ruscetti et al., 2016). More importantly, these models motivated investigating the behavior of a set of non-small cell lung cancer (NSCLC) cell lines categorized as ‘hybrid’ based on bulk measurements (Schliekelman et al., 2015) at the single-cell level. The ‘hybrid’ cell lines contained subpopulations of epithelial and mesenchymal cells (H2291) (Figure 2B) and/or individual cells co-expressing epithelial and mesenchymal markers, such as CDH1 and VIM (H1975). H1975 cells exhibited a hybrid E/M phenotype at the single-cell level over multiple passages (Figure 3A), strongly suggesting that a hybrid E/M state can be a stable phenotype (Jolly et al., 2016). As compared to epithelial cells (H820), H1975 cells also stained for nuclear ZEB1 (Jia et al., 2017), thus confirming the prediction made by the mathematical model developed by Lu *et al*. (Lu et al., 2013) (Figure 3B).

**Figure 2:**
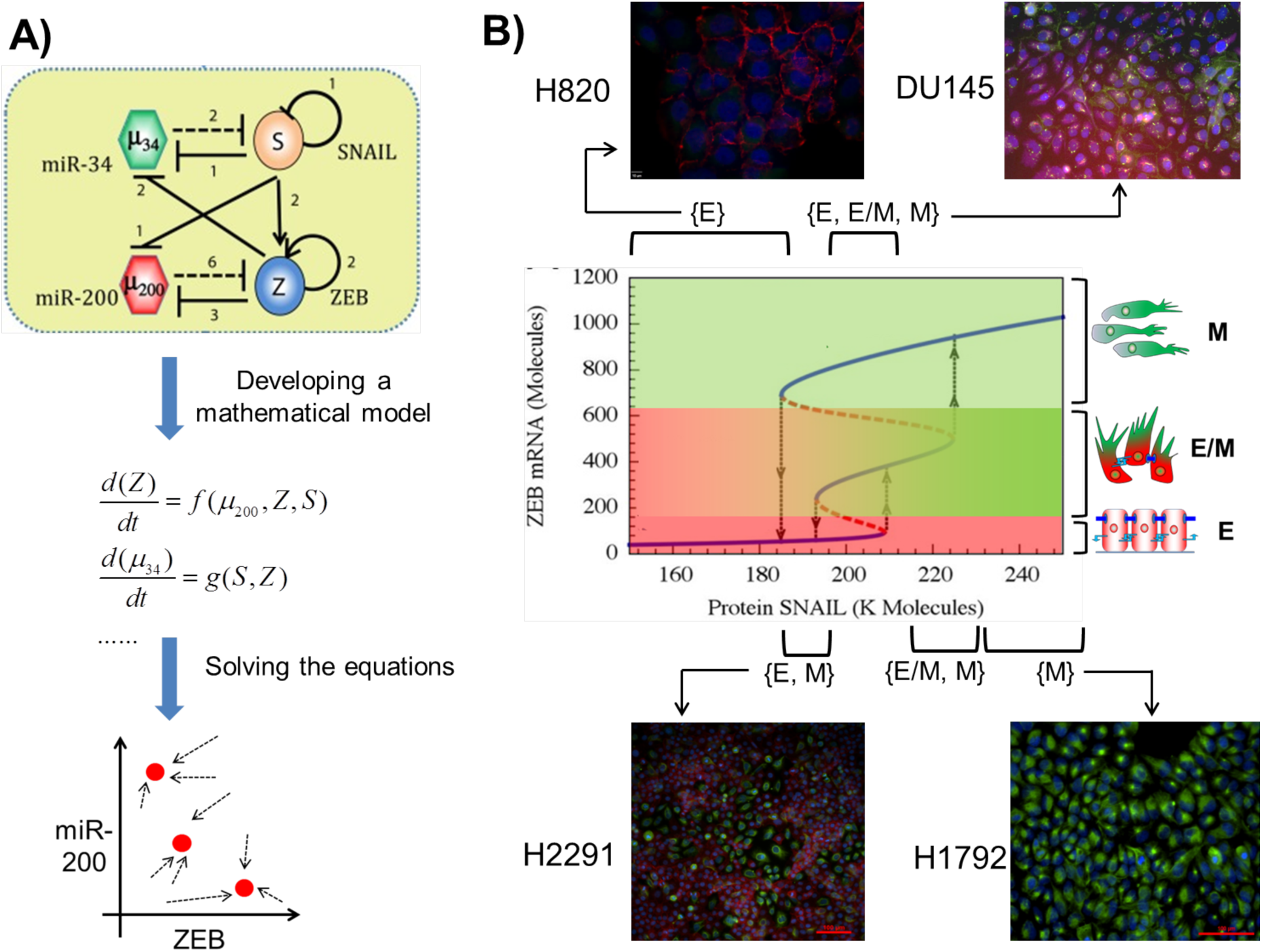
Integrated theoretical-experimental framework to understand epithelial-mesenchymal plasticity. **A**) (Top) EMT regulatory circuit denoting two transcription factor families – SNAIL and ZEB, and two microRNA (miR) families – miR-34, miR-200. Transcriptional (denoted by solid lines) and miR-mediated (denoted by dotted lines) regulations in this circuit can be represented as a set of mathematical equations (middle) that can then be solved to attain the steady states or phenotypes (shown by red solid dots) and dynamics of this circuit. **B**) (middle) Bifurcation diagram depicting the change in ZEB mRNA levels, and consequently phenotypic switching (shown by black arrows), for varying values of SNAIL. Solid blue lines depict stable states (phenotypes), and dotted red lines illustrate unstable states. Mesenchymal cells have highest levels of ZEB mRNA (topmost blue line), followed by hybrid E/M cells (middle blue line) and then epithelial cells (blue line at the bottom). (Top and bottom) Immunofluorescence staining for CDH1 (red) and VIM (green) in different cancer cell lines reveal the existence of individual phenotypes or co-existence of more than one phenotypes, as predicted by the mathematical model. Cell lines corresponding to each region are marked; for instance, H2291 cell populations contains cells staining for either CDH1 or VIM, but not individual cells co-staining for CDH1 or VIM, thus H2291 maps on to the region where cells can adopt either an E or a M state – {E, M}.

Observations in H1975 serve as a remarkable example of the power of leveraging an integrated theoretical-experimental framework. Although the mathematical models predicted regions for a coexistence of {E, E/M, M} and {E/M, M} states, a parameter region enabling a hybrid E/M state alone, i.e. {E/M}, predominantly was not observed (Figure 2). Consequently, that led to a search for potential ‘phenotypic stability factors’ (PSFs) – molecular players that can enable a monostable {E/M} region. Incorporating two proteins OVOL2 and GRHL2 that were reported to form mutually inhibitory loops with ZEB (Cieply et al., 2013, 2012; Roca et al., 2013) – in the mathematical model predicted the existence of a desired {E/M} region (Hong et al., 2015; Jia et al., 2015; Jolly et al., 2016) (Figure 3A). The role of OVOL2 and GRHL2 as PSFs was validated by experiments showing that knockdown of either of these proteins in H1975 drove the cells toward a stable hybrid E/M state to a fully mesenchymal phenotype (Jolly et al., 2016). Similar results in developmental EMT contexts strengthened the notion that these PSFs can act as ‘molecular brakes’ on EMT that can prevent cells ‘that have gained partial plasticity’ from undergoing a complete EMT (Watanabe et al., 2014; Werner et al., 2013). Furthermore, the mathematical model suggested that overexpression of PSFs can drive an MET, a prediction already verified in breast and prostate cancer cell lines (Roca et al., 2013; Werner et al., 2013), and kidney cells (Aue et al., 2015), thereby indicating that such models can behave as ‘semi-quantitative predictive paradigms’ to predict the cellular behavior pertinent to EMT regulation in multiple cell lines.

**Figure 3:**
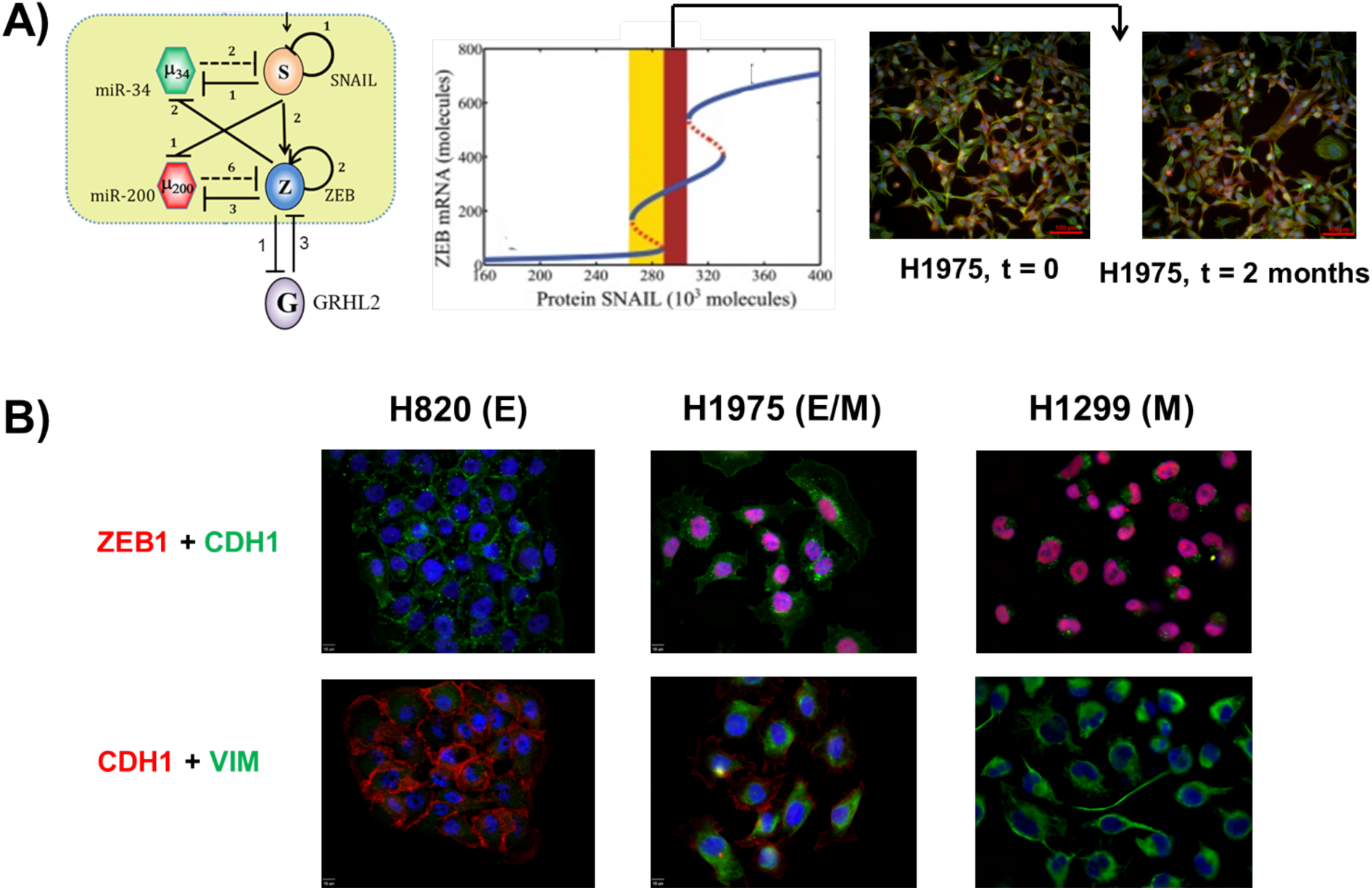
Characterizing a hybrid epithelial/mesenchymal (E/M) phenotype. **A**) (left) EMT circuit as shown earlier, with GRHL2 being incorporated based on literature about its interactions with ZEB. (middle) Bifurcation diagram depicting change in the levels of ZEB mRNA as a function of varying SNAIL levels, corresponding to the circuit diagram shown in left. It illustrates a monostable {E/M} region highlighted by brown color. (right) Immunofluorescence images for CDH1 and VIM for H1975 over multiple passages consistently shows single-cell co-expression for both markers. **B**)

Despite the utility of these models, it is important to note that we neither claim that these particular models can accurately predict EMT regulation for all cell lines nor that they can necessarily predict responses to all perturbations that can alter EMT status in a given cell line. For instance, overexpression of GRHL2 did not drive MET in the RD and 143B human sarcoma cell lines (Somarelli et al., 2016). Further experiments indicated that in RD and 143B, GRHL2 coupled to miR-200 and ZEB1 in a different topology as compared to that in multiple (adeno) carcinoma cell lines. In sarcoma cells, GRHL2 had no effect on ZEB1 levels, and GRHL2-induced changes were only observed when ZEB1 was knocked down (Figure 3). These findings led to the development of a revised mathematical model that captured these newly-revealed interactions. The revised model was able to reproduce robustly the key features of experiments in RD and 143B, such as the synergistic induction of E-cadherin levels upon overexpression of both GRHL2 and miR-200 (Somarelli et al., 2016), and predicted how epigenetic regulation of GRHL2 can modulate MET. Therefore, ‘no one size fits all’; no model – either biological or mathematical – fits all different biological contexts; carcinoma cell lines may not be reliable biological models to understand sarcoma biology, and similarly networks that work well for predicting carcinoma cell line behavior need not be the same for sarcoma ones.

Notwithstanding the complexity and heterogeneity in the gene regulatory networks that drive EMT and MET in different contexts, mathematical models can be constructed to help rationalize existing experimental results and to aid in guiding further experiments, by making certain approximations or estimations about the model parameters. Each of the mathematical models developed above has multiple variables – ZEB, miR-200, GRHL2 etc. – and each variable is represented by an equation tracing their levels over time. Each equation has terms representing the innate production and degradation rates for those species that can be estimated from their half-lives and/or typical number of molecules in a cell (Milo et al., 2010). Similarly, each equation contains terms pertaining to regulation of the respective species by one another; for instance, inhibition of ZEB by miR-200. The quantitative parameters describing these interactions, such as the number of binding sites and the fold-change in levels upon overexpression or inhibition, can also be gained from relevant experimental data. For example, whereas miR-200s can bind up to eight to nine binding sites on Zeb mRNA and reduces the protein levels by 90% (Gregory et al., 2011), miR-34 binds to two binding sites on Snail mRNA and reduces the protein levels only by 50% (Kim et al., 2011). Upon estimating a relevant range of parameter variation, the sensitivity of these models to different parameters can be tested. For instance, the range of levels of SNAIL for which a hybrid E/M phenotype is observed is largely robust to ± 20% variation in parameters (Jia et al., 2015). Thus, one need not know the exact value of each parameter in the mathematical model for every cell line. Instead, estimating their typical range from the experimental data can be a good first approximation. This approximation is good because it can be often impossible to perform all experiments to measure every single parameter for every single cell line, and these measurements can themselves be subject to uncertainty (Azeloglu and Iyengar, 2015; Kirk et al., 2015).

Deriving mathematical models to represent biological systems is rarely straightforward (Kirk et al., 2015). Thus a key to justifiably use mathematical models is to state the assumptions and uncertainty in the model structure and/or parameters clearly. If one believes the assumptions of the model, one must also believe its conclusions (Gunawardena, 2014) – and this applies both to mathematical and biological models. For instance, in models of the EMT/MET regulatory network described above (Lu et al., 2013; Tian et al., 2013), more than one family member of a protein or microRNA are lumped into one variable, for the sake of simplicity. So, an implicit assumption of these mathematical models is that, for instance, both ZEB1 and ZEB2 behave identically, which need not be true in all contexts. Similarly, in the context of biological models, an underlying assumption in *in vitro* cell culture is that the observed behavior of cells in a two-dimensional setup plated on plastic recapitulates the ‘true’ behavior of cells *in vivo*.

### How can mathematical models be used to study changes in other cellular traits connected with EMT/MET?

EMT and MET are considered as the motors of cellular plasticity due to their coupling with other cellular traits such as metabolism, tumor-initiation potential, genome plasticity, drug resistance, immune-suppression, cell-cell communication etc. (Brabletz et al., 2011; Chen et al., 2014; Fischer et al., 2015; Lu et al., 2014; Mani et al., 2008; Morel et al., 2008; Tripathi et al., 2016; Wellner et al., 2010; Zheng et al., 2015). By using mathematical models similar to those described above, one can investigate the interplay of EMT/MET with any one or more of these traits.

For instance, mathematical models have helped reconcile apparently contradictory results with regards to the interplay between EMT/MET and ‘stemness’ or tumor-initiation potential. Initially, EMT was proposed to promote a gain of stem-like properties (Mani et al., 2008; Morel et al., 2008). However, later studies suggested that cells locked in a mesenchymal phenotype often lose their stem-like traits (Celià-Terrassa et al., 2012; Tran et al., 2014), and that both epithelial-like and mesenchymal-like stem-like subpopulations may exist (Liu et al., 2014) (Figure 4A, i-iii). To provide a unifying schema to explain these apparently conflicting results, a mathematical model was developed to connect core EMT players, miR-200 and ZEB, with the master regulators of stemness, LIN28 and let-7 (Yang et al., 2010). This model proposed that cells in a hybrid E/M phenotype can be more likely to gain stemness as compared to those in either a fully epithelial or mesenchymal state (Jolly et al., 2014) (Figure 4B, i-ii). Follow-up experiments in breast cancer cells demonstrated that hybrid E/M cells – cells co-expressing canonical epithelial and mesenchymal genes to a similar level – can form up to 10-times more mammospheres as compared to strongly epithelial or mesenchymal cells, thus validating the prediction of the model (Grosse-Wilde et al., 2015) (Figure 4B,iii). Hybrid E/M cells also drove aggressive tumor growth *in vivo* (Goldman et al., 2015). Moreover, enhanced or acquired drug resistance of breast cancer and oral squamous carcinoma cells in a hybrid E/M phenotype further substantiate the proposed correlation between a hybrid E/M phenotype and ‘stemness’ (Biddle et al., 2016; Brown et al., 2016; Goldman et al., 2015). Despite initial promising validations, further research is needed to evaluate how well the hypothesis holds that the hybrid E/M state is more stem-like (Celià-Terrassa and Kang, 2016). Moreover, the positioning of a ‘stemness window’ need not be fixed mid-way on the EMT axis, but could instead be much more dynamic and subtype- and/or patient-specific.

**Figure 4:**
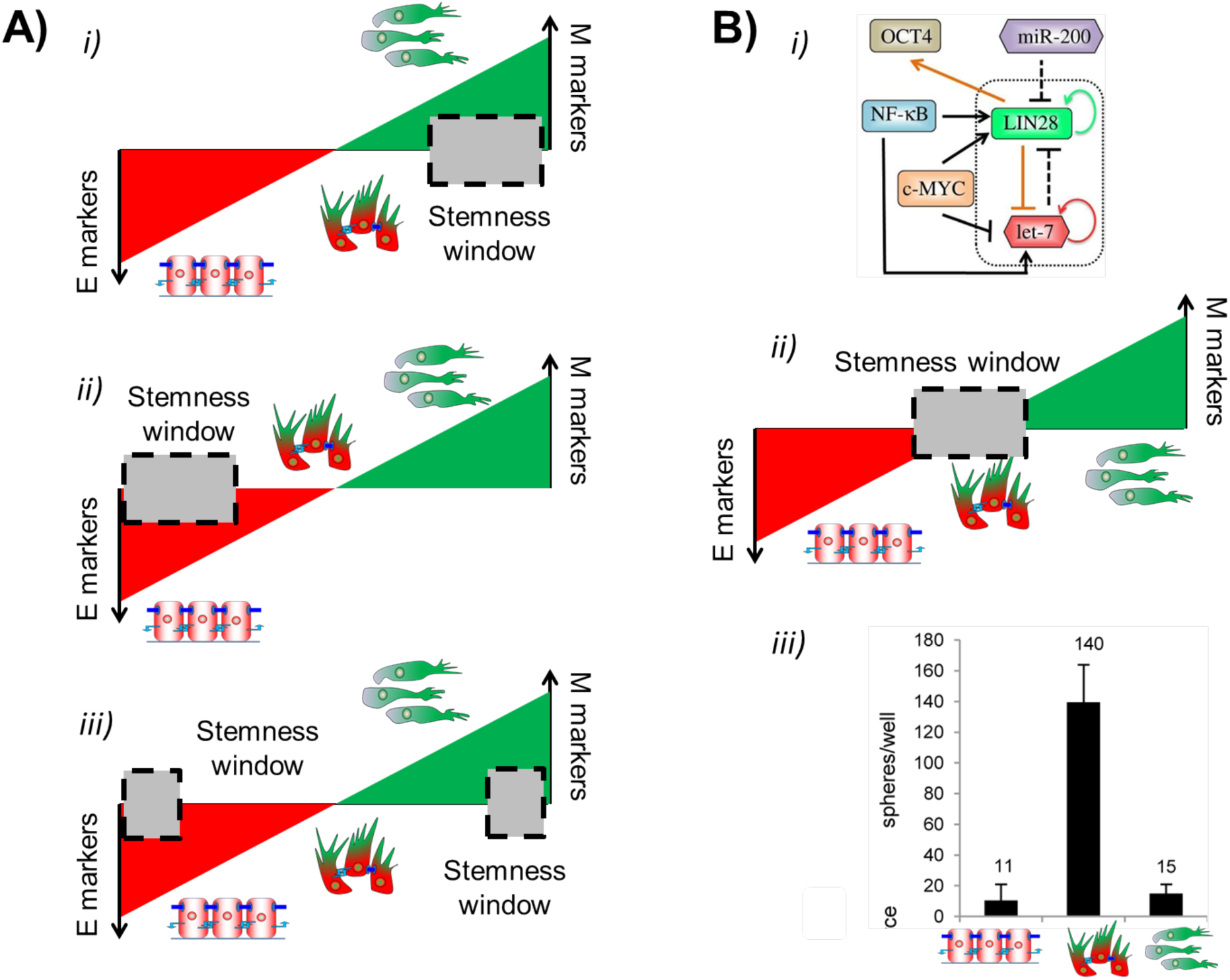
EMT-stemness interplay. **A**) Schematics representing apparently contradictory results on the EMT status of cancer stem cells (CSCs) (left), as shown by the position of ‘stemness window’ on the ‘EMT axis’ with E and M as two ends. **B**) (top) Circuit simulated via mathematical model by Jolly et al. J R Soc Interface 2014 for decoding EMT-stemness interplay. (middle) Prediction of the mathematical model about the location of a ‘stemness window’. (bottom) Experiments showing the relative tumor-initiation potential of E, hybrid E/M and M subpopulations (Modified from Grosse-Wilde et al. PLoS ONE 2015).

Similarly, in a study demonstrating that a mesenchymal phenotype correlates with immune evasion via reduced expression of the immunoproteasome, a mathematical model was developed to capture an underlying mechanism of immunoproteasome regulation that involved STAT3, STAT1 and miR-200s (Tripathi et al., 2016). The model predicted that inhibiting the activation of STAT3 can increase the levels of immuno-proteasome subunits PSMB8 and PSMB9 in mesenchymal NSCLC cell lines. Indeed, inhibition of STAT3 using rapamycin led to enhanced levels of PSMB8 and PSMB9 via an activated STAT1 pathway.

Another specific question where mathematical models may prove to be crucial to decode the underlying dynamics is the epigenetic reprogramming accompanying EMT/MET (Tam and Weinberg, 2013). The ‘poised’ chromatin state of ZEB1 in which the ZEB1 promoter simultaneously displays epigenetic marks of both active and repressed chromatin may enhance cellular plasticity among cancer stem cells (CSCs) and non-CSCs and consequently spike tumorigenic potential (Chaffer et al., 2013). Similarly, epigenetic differences can modulate MET induction in sarcomas (Somarelli et al., 2016). Finally, these epigenetic interactions could possibly modulate the transition rates among epithelial, mesenchymal and hybrid E/M phenotypes in specific cell lines by controlling genome-wide chromatin marks. A quantitative comparison of transition rates as measured using various reporter systems (Somarelli et al., 2013; Toneff et al., 2016) and those predicted by modeling of the underlying regulatory networks (Li et al., 2016) can bridge the gaps in our understanding of epithelial-mesenchymal plasticity. Similarly, existing theoretical frameworks to investigate epigenetic regulation (Steffen et al., 2012) can be integrated with mathematical models incorporating interconversion among CSCs and non-CSCs (Li and Wang, 2015; Yang et al., 2012; Zhou et al., 2013) and temporal mapping of epigenetic changes during EMT/MET (Kao et al., 2016) to identify the epigenetic marks that can be targeted to constrain cellular plasticity and thus abate metastatic and therapy-resistant progression.

### How can mathematical models connect signaling aspects to cellular motility associated with EMT/MET?

Altered cellular motility and cellular morphology traits are considered to be the primary biophysical consequence of EMT/MET. During EMT, cells typically have reduced adhesion with their neighbors, and migrate collectively or individually depending on their intercellular adhesion and spatial confinement (Boekhorst et al., 2016). For instance, during embryonic development, neural crest cells undergoing a partial or complete EMT can migrate as either a multicellular stream or individually, in order to reach distant tissues. Similarly, during gastrulation, both these modes of migration are observed at different spatiotemporal coordinates (Scarpa and Mayor, 2016). Typically, collective migration is associated with a partial EMT or hybrid E/M phenotype (Kuriyama et al., 2014; Sarioglu et al., 2015), whereas fully mesenchymal cells tend to migrate alone. Depending on cell-matrix adhesion, the migrating cells can also reversibly switch to an amoeboid migration mode, where cells migrate individually and predominantly via squeezing through the gaps in extracellular matrix (ECM) (Pankova et al., 2010; Wolf et al., 2007). Similar to EMT/MET, the choice between mesenchymal and amoeboid modalities need not be a binary process and cells can exhibit signatures of both mesenchymal and amoeboid motility - lamellopodia and bleb-like protrusions respectively (Bergert et al., 2012; Yoshida and Soldati, 2006). Preliminary mathematical models of some of the underlying signaling mechanisms governing these transitions have been developed (Huang et al., 2015, 2014), but a detailed analysis of how these molecules impinge upon changes in cytoskeletal reorganization, cell shape, cell-cell adhesion, cellular contractility, and cell-ECM mechanics and consequently drive different migration modes remains to be accomplished.

Multiple existing theoretical approaches for cell motility models focus on these key mechanical aspects. Most frameworks for single-cell migration have focused on and driven further experiments in fish keratocytes (Holmes and Edelstein-Keshet, 2012; Ziebert et al., 2012). For instance, Shao *et al*. (Shao et al., 2012) illustrate how cell morphology is determined by collective effects of myosin contraction, actin polymerization, and adhesion site dynamics. This type of approach could actually make contact with the time-course data correlating cell shape to EMT states (Mandal et al., 2016; Sarkar et al., 2016). In contrast to these single-cell models, other frameworks have concentrated on tissue-level dynamics by constructing models for adhesive cell clusters and monolayers (Basan et al., 2013; Bi et al., 2016; Harris et al., 2012; Kabla, 2012; Zimmermann et al., 2016) (Figure 5A, B). In addition to actomyosin dynamics, these models can incorporate intercellular forces, cell density, substrate properties, and contact inhibition of locomotion (CIL) – a fundamental feature of collective cell migration that promotes the formation of protrusions in a direction away from their contacts with the follower cells, thereby propelling the migration by leader cells (Figure 5C) (Mayor and Etienne-Manneville, 2016). With an emerging understanding of mechanochemical coupling regulating the determination of leader and follower cells (Riahi et al., 2015), the models described above focusing on tissue dynamics can elucidate how different signaling aspects crosstalk with cell and tissue mechanics during collective cell migration.

**Figure 5:**
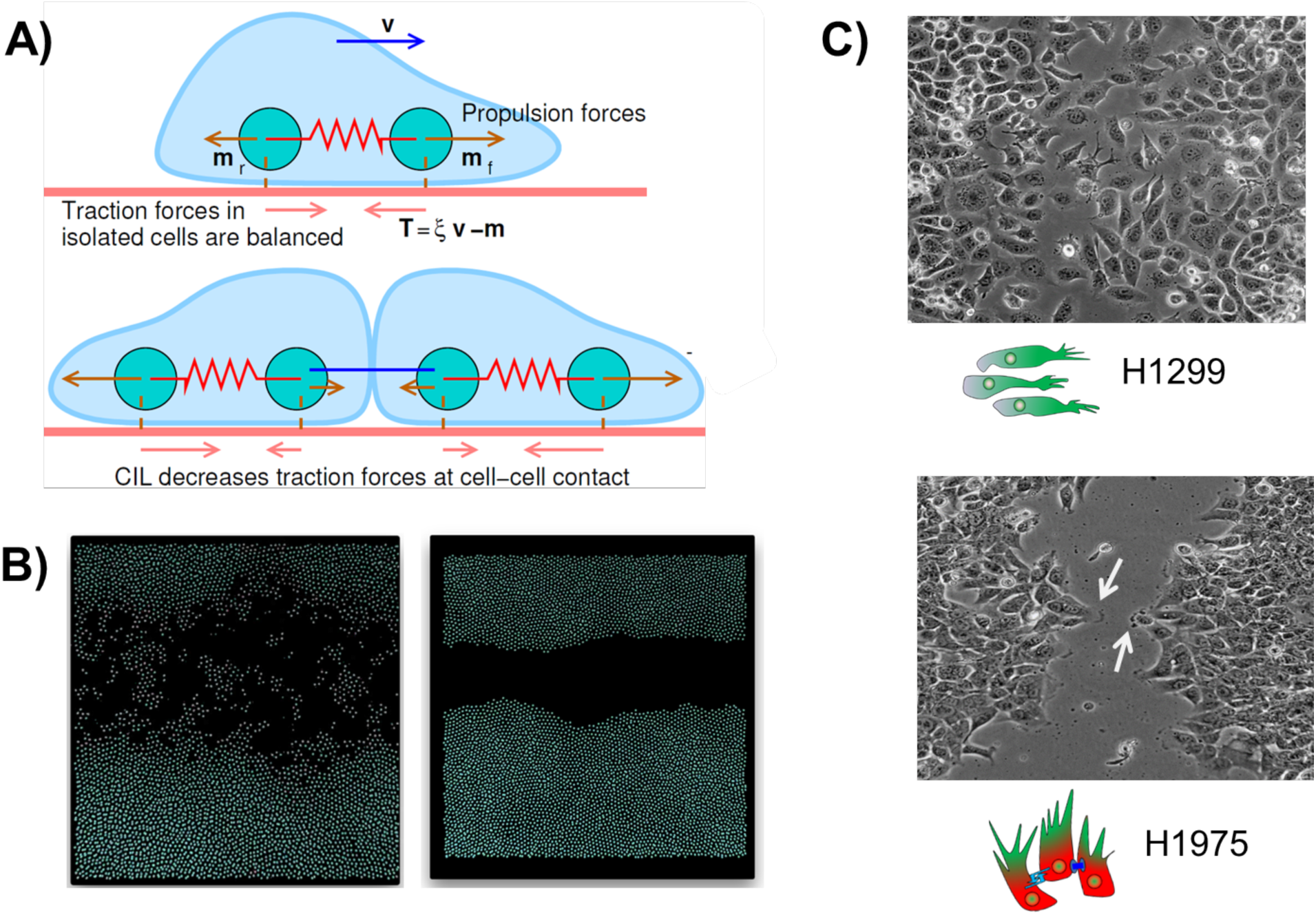
Mathematical models for cell motility. **A**) Each cell is represented by two particles, both of which exert forces on the substrate. Upon cell-cell contact, due to contact inhibition of locomotion, these forces change in magnitude and direction. (Reprinted from Zimmermann et al. PNAS 2016) **B**) Simulations for individual cell migration (left) and collective cell migration (right); shown is one snapshot emerging from this model of cell motility. **C**) Individual migration observed for mesenchymal cell line H1299 and collective migration with the emergence of leader cells (highlighted by arrow) forming finger-like projections observed for H1975 (hybrid E/M cell line) – reproduced from Jolly et al. Oncotarget 2016

In terms of its application to cancer, a form of collective cell migration where multicellular clusters of tumor cells can bud off the primary lesions and enter circulation, has been observed to be the predominant way of successful colonization (Aceto et al., 2014; Cheung et al., 2016). These clusters of Circulating Tumor Cells (CTCs) retain their epithelial traits, at least partially, and act as primary harbingers of metastasis (Cheung and Ewald, 2016; Grigore et al., 2016; Jolly et al., 2015). Differential gene expression signatures of leader vs. follower cells in collective migration and invasion during metastasis has highlighted JAG1 as a key player (Cheung et al., 2016), thereby reminiscent of the involvement of Notch signaling in regulating leader vs. follower phenotypes in multiple contexts of collective cell migration (Blanco and Gerhardt, 2013; Boareto et al., 2015; Riahi et al., 2015).

This connection between Notch signaling and collective migration motivated a recently developed mathematical model that incorporated the coupling between EMT circuit and Notch signaling pathway as based on experimental data (Brabletz et al., 2011; Bu et al., 2013; de Antonellis et al., 2011; Niessen et al., 2008; Sahlgren et al., 2008). This model predicted that Notch-Jagged signaling, but not Notch-Delta signaling, can enable both increased numbers and spatial proximity of hybrid E/M cells that, owing to their ability to both adhere and migrate, may lead to formation of clusters of CTCs (Boareto et al., 2016). This prediction provides mechanistic insights into why JAG1 may be crucial for mediating clustered migration (Cheung et al., 2016), and is consistent with the evidence that JAG1 is related to drug resistance (Boareto et al., 2016; Guo et al., 2010), if we refer to the earlier claim that hybrid E/M cells are more likely to exhibit stemness. Yet, it remains to be rigorously and extensively tested experimentally whether knockdown of JAG1 can reduce the frequency of clustered migration and thereby curtail metastasis.

For a comprehensive characterization of collective cell migration in cancer, such signaling mechanism-based models need to be tied to previously described models of cell motility in multiple ways, for instance, by incorporating the effect of cellular stress on the activation of Notch signaling (Riahi et al., 2015), integrating how matrix stiffness can drive EMT through TWIST1-GP3B2 pathway (Wei et al., 2015), including how matrix density can alter the levels of membranous E-cadherin and affect the EMT status of cells (Kumar et al., 2014), and considering ZEB1 induces collagen deposition and stabilization (Peng et al., 2016). Developing such mechanochemical models offers can reveal how phenotypic transitions are coupled to the repertoire of mechanical signals that cancer cells experience and generate (Przybyla et al., 2016).

### What other open questions in the regulation of EMT/MET can benefit from mathematical models?

Multiple open questions related to EMT/MET furnish exciting opportunities for cross-pollination of ideas among experimental and computational biologists, including (a) how many intermediate states can cells attain *en route* to EMT and MET?; (b) what is the genomic, proteomic, and epigenetic signature of these states?; (c) how symmetric are the dynamics of EMT and MET, and whether cells display hysteresis (i.e. cellular memory)?; and (d) what is the relative stability and relative ‘stemness’ possessed by each of these states? As expected, mathematical models encompassing a larger number of EMT/MET regulatory players than considered in the initial models (Lu et al., 2013; Tian et al., 2013) have suggested multiple intermediate states (Hong et al., 2015; Huang et al., 2016; Steinway et al., 2015), but these predictions remain to be experimentally verified, thus providing impetus for many collaborative efforts.

Further, epithelial-mesenchymal plasticity of cancer cells has also been linked to metabolic shifts (Dong et al., 2013; Kondaveeti et al., 2015; LeBleu et al., 2014) – another hallmark of cancer (Hanahan and Weinberg, 2011). Mathematical models which calculate metabolic fluxes by considering mass balance of various intra-cellular metabolites is a standard technique to analyze metabolic signatures (Markert and Vazquez, 2015). Such models are being increasingly implemented to quantify metabolic changes in tumor cells (Achreja et al., 2017). Constructing mathematical modeling frameworks that integrate these flux-balance models with models for the dynamics of signaling networks can help investigate the coupling of metabolic networks with signaling pathways regulating epithelial-mesenchymal plasticity and stemness (Menendez and Alarcón, 2014; Peiris-pagès et al., 2016) can offer novel insights into the emergent consequences of bidirectional crosstalk among these networks driving these different hallmarks of cancer.

In addition to discerning this intracellular crosstalk, mathematical models can infer the dynamics of stromal cells as well as intercellular tumor-stroma signaling and act as *in silico* co-culture systems. For instance, mechanism-based mathematical models can explain how macrophages can exhibit an intermediate polarization status between M1 and M2 (Italiani and Boraschi, 2014). Further, models capturing the crosstalk between differentially polarized macrophages and cancer cells (Yang et al., 2016) at an intra-cellular decision-making level as well as at a population level (i.e. multi-scale models) can help visualize how cancer cells can engineer their microenvironment to their benefit and drive tumor progression, and hence propose strategies to restrict it.

## Conclusion

As discussed above, an integrated theoretical-experimental approach has been instrumental in characterizing epithelial-mesenchymal plasticity and cellular traits associated with this plasticity. Concomitant with the renewed understanding that cancer can be viewed as an ecosystem unto itself (Yang et al., 2014), mathematical models capturing the interplay between tumor cells and multiple components of the tumor microenvironment can decode underlying organizing principles that manifest as myriad phenotypic complexities (Hanahan and Weinberg, 2011). Therefore, an iterative crosstalk between theory and experiment can help propel the hope that cancer biology and treatment ‘ will become a science with a conceptual structure and logical coherence that rivals that of chemistry or physics’ (Hanahan and Weinberg, 2000) into reality.

## Acknowledgements

This work was supported by the National Science Foundation (NSF) Center for Theoretical Biological Physics (NSF PHY-1427654) and NSF DMS-1361411. HL was also supported as a CPRIT (Cancer Prevention and Research Institute of Texas) Scholar in Cancer Research of the State of Texas at Rice University. SH was supported by the Rubenstein Family Foundation and the Canary Foundation. JAS acknowledges support from the Duke Cancer Institute, the Duke Genitourinary Oncology Laboratory, the Department of Orthopaedics, and the Triangle Center for Evolutionary Medicine (TriCEM)

